# Stochastic fluctuations of the facultative endosymbiont *Wolbachia* due to finite host population size

**DOI:** 10.1101/2025.02.19.638912

**Authors:** Jason M. Graham, Joseph Klobusicky, Michael T.J. Hague

## Abstract

Many insects and other animals host heritable endosymbionts that alter host fitness and reproduction. The prevalence of facultative endosymbionts can fluctuate in host populations across time and geography for reasons that are poorly understood. This is particularly true for maternally transmitted *Wolbachia* bacteria, which infect roughly half of all insect species. For instance, the frequencies of several *w*Mel-like *Wolbachia*, including *w*Mel in host *Drosophila melanogaster*, fluctuate over time in certain host populations, but the specific conditions that generate temporal variation in *Wolbachia* prevalence are unresolved. We implemented a discrete generation model in the new R package *symbiontmodeler* to evaluate how finite-population stochasticity contributes to *Wolbachia* fluctuations over time in simulated host populations under a variety of conditions. Using empirical estimates from natural *Wolbachia*-*Drosophila* systems, we explored how stochasticity is determined by a broad range of factors, including host population size, maternal transmission rates, and *Wolbachia* effects on host fitness (modeled as fecundity) and reproduction (cytoplasmic incompatibility; CI). While stochasticity generally increases when host fitness benefits and CI are relaxed, we found that a decline in the maternal transmission rate had the strongest relative impact on increasing the size of fluctuations. We infer that non- or weak-CI-causing strains like *w*Mel, which often show evidence of imperfect maternal transmission, tend to generate larger stochastic fluctuations compared to strains that cause strong CI, like *w*Ri in *D. simulans*. Additional factors, such as fluctuating host fitness effects, are required to explain the largest examples of temporal variation in *Wolbachia*. The conditions we simulate here using *symbiontmodeler* serve as a jumping off point for understanding drivers of temporal and spatial variation in the prevalence of *Wolbachia*, the most common endosymbionts found in nature.

## INTRODUCTION

Animal life is characterized by diverse interactions with microbes, with relationships spanning a continuum from beneficial to antagonistic. Many insects and other animals carry heritable endosymbionts that are vertically-maternally transmitted from one host generation to the next (Russell et al. 2019; Hague et al. 2024). Endosymbionts can alter basic aspects of host fitness, including reproduction, immune function, and nutrient acquisition (McFall-Ngai et al. 2013; Shropshire et al. 2020; Perreau and Moran 2022; Bennett et al. 2024; Hoffmann and Cooper 2024). The fitness consequences for hosts are ultimately determined by whether or not endosymbionts are prevalent in host populations. Many endosymbionts are facultative from the host perspective, such that a proportion of host individuals in the population do not carry endosymbionts (Mateos et al. 2006; Carrington et al. 2011; Hamm et al. 2014; Corbin et al. 2017; Hague et al. 2020*b*). The frequency of facultative endosymbionts in host populations can vary widely across time, geography, and host systems for reasons that are poorly understood (Hamm et al. 2014; Kriesner et al. 2016; Cooper et al. 2017; Hague et al. 2020*b*; Smith et al. 2021; Wheeler et al. 2021; Gimmi et al. 2023; McPherson et al. 2023).

Maternally transmitted *Wolbachia* are the most common endosymbionts on earth, infecting roughly half of all insect species, as well as other arthropods and nematodes (Ferri et al. 2011; Zug and Hammerstein 2012; Weinert et al. 2015). *Wolbachia* are maternally transmitted through the female germline (Russell et al. 2019; Porter and Sullivan 2023; Radousky et al. 2023; Hague et al. 2024); although, horizontal and introgressive transfers are common on evolutionary timescales (O’Neill et al. 1992; Raychoudhury et al. 2009; Conner et al. 2017; Gerth and Bleidorn 2017; Turelli et al. 2018; Cooper et al. 2019; Vancaester and Blaxter 2023; Shropshire et al. 2024). Patterns of *Wolbachia* spread in nature indicate that the endosymbionts are generally beneficial for host fitness, although these effects are poorly understood in natural populations (Hoffmann and Turelli 1997; Kriesner et al. 2013, 2016; Hamm et al. 2014; Kriesner and Hoffmann 2018; Meany et al. 2019). Recent work suggests *Wolbachia* block viruses in their native *Drosophila* hosts (Hedges et al. 2008; Teixeira et al. 2008; Osborne et al. 2009; Martinez et al. 2014; Cogni et al. 2021; Bruner-Montero and Jiggins 2023), in addition to a potential role of nutrient provisioning (Brownlie et al. 2009; Nikoh et al. 2014; Newton and Rice 2020). Some *Wolbachia* strains also cause cytoplasmic incompatibility (CI), a crossing incompatibility that generates a frequency dependent benefit that favors *Wolbachia*-positive females (Shropshire et al. 2020; Turelli et al. 2022).

*Wolbachia* typically occur as facultative endosymbionts in insects (from the host perspective) and are only found in a proportion of host individuals within a given population. *Wolbachia* frequencies can vary considerably in host populations across time and geography for reasons that are poorly understood (Hoffmann et al. 1990; Kriesner et al. 2013, 2016; Hamm et al. 2014; Cooper et al. 2017; Hague et al. 2020*b*, 2022; Wheeler et al. 2021; Turelli et al. 2022). This seems to be particularly true for *Wolbachia* strains that cause no or weak CI. For example, the *w*Mel strain in *D. melanogaster* has been shown to fluctuate dramatically over time at a single locale, with frequency fluctuations >0.7 in some instances and no evidence of seasonality (Hoffmann et al. 1998; Reynolds and Hoffmann 2002). Smaller temporal fluctuations have been observed for the related “*w*Mel-like” *w*San and *w*Yak strains (diverged from *w*Mel ∼30,000 years ago) that cause weak CI in *D. yakuba*-clade hosts on the island of São Tomé off the coast of west Africa (Cooper et al. 2017, 2019; Hague et al. 2020*b*). These fluctuating patterns contrast those of strong CI-causing *Wolbachia* strains (e.g., *w*Ri in *D. simulans*), which tend to persist at high, stable frequencies (Hoffmann et al. 1990; Turelli and Hoffmann 1991, 1995; Rousset and Solignac 1995; Bourtzis et al. 1996; James and Ballard 2000; Ballard 2004; Carrington et al. 2011; Choi et al. 2015; Turelli et al. 2018). Examples of temporal frequency variation for strains like *w*Mel suggest that *Wolbachia* may fluctuate stochastically in host populations under certain conditions (Jansen et al. 2008; Kriesner and Hoffmann 2018; Turelli and Barton 2022; Turelli et al. 2022).

We used mathematical models to evaluate how finite-host population stochasticity contributes to *Wolbachia* fluctuations over time in simulated host populations. Population frequencies and equilibria dynamics of *Wolbachia* can be approximated by a discrete generation, deterministic model that incorporates three parameters: (1) the proportion of uninfected ova produced by *Wolbachia*-positive females (*µ*; i.e., imperfect maternal transmission), (2) the fitness of *Wolbachia*-positive females relative to *Wolbachia*-negative females (*F*; i.e., components of host fitness like fecundity), and (3) the reduction in the relative egg hatch of uninfected eggs fertilized by *Wolbachia*-positive males due to CI (*s_h_*; i.e., CI strength) (Figure 1) (Hoffmann et al. 1990). Following Turelli et al. (2022) and Turelli and Barton (2022), we adapted the deterministic model of *Wolbachia* equilibria to approximate the stochasticity induced by finite population size using a stochastic transition matrix analogous to a haploid Wright-Fisher model of genetic drift (Crow and Kimura 1970; Turelli and Barton 2022; Turelli et al. 2022). The model uses an effective population size of *N* female hosts (because *Wolbachia* are maternally transmitted) and binomial sampling to model the stochastic effects of finite population size in the context of the maternal transmission rate (*µ*), host fitness effects (*F*), and the strength of CI (*s_h_*).

**Figure 1.**
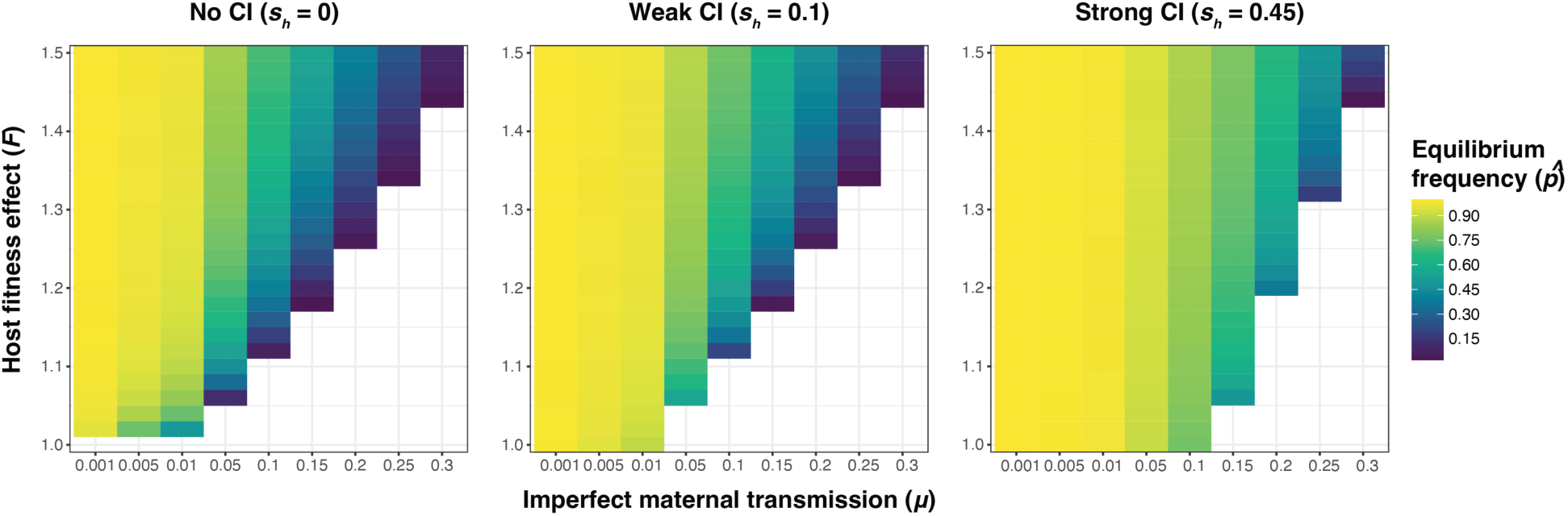
Equilibrium *Wolbachia* frequencies (*p*^) approximated by a discrete generation deterministic model incorporating imperfect maternal transmission (*µ*), *Wolbachia* effects on host fitness (*F*), and cytoplasmic incompatibility (*s_h_*). Equilibrium frequencies are shown across a range of biologically plausible parameter values for no, weak, and strong CI.

We implemented the simulations in a new publicly available R package called *symbiontmodeler*. We used the R package to evaluate how biologically plausible values of host population size (10^3^ ≤ *N* ≤ 10^6^), imperfect maternal transmission (0.001 ≤ *µ* ≤ 0.3), host fitness effects (1 ≤ *F* ≤ 1.5), and CI (*s_h_* = 0, 0.1, 0.45) influence the size of stochastic *Wolbachia* fluctuations over time in simulated host populations. The package includes additional features to consider a variety of conditions that may occur in natural host populations. For instance, the binomial *µ* parameter represents imperfect maternal transmission and we allow for inclusion of a subpopulation of “low transmitter” females with especially poor transmission rates, because field estimates of maternal transmission for *w*Mel-like *Wolbachia* suggest that a small proportion of females may have very low transmission (i.e., *µ* > 0.7) (Hoffmann et al. 1998; Carrington et al. 2011; Hamm et al. 2014; Hague et al. 2020*b*). We also allow for the option of fluctuating host fitness effects by treating *F* as a log-normal random variable (Turelli et al. 2022), because some evidence suggests that the fitness effects of *w*Mel may be context-dependent (Kriesner et al. 2016; Chrostek et al. 2021). After simulating a wide range of conditions, our results demonstrate how non-CI-causing strains like *w*Mel, which show evidence of imperfect maternal transmission, tend to generate larger stochastic fluctuations over time, as compared to strong-CI-causing strains like *w*Ri.

## THEORETICAL FRAMEWORK

### Deterministic analysis of Wolbachia frequencies

The deterministic model of *Wolbachia* population frequencies and equilibria dynamics incorporates imperfect maternal transmission (*µ*), host fitness effects modeled as differential fecundity (*F*), and CI strength (*s_h_*) (Hoffmann et al. 1990). We assume that uninfected ova produced by *Wolbachia*-positive females (due to imperfect maternal transmission) are susceptible to CI, just as uninfected ova from *Wolbachia*-negative females (Carrington et al. 2011). Embryos produced by fertilizations of *Wolbachia*-negative ova with sperm from *Wolbachia*-positive males hatch with frequency *H* = 1 – *s_h_* relative to the other three possible fertilizations, which are all considered equally compatible (Cooper et al. 2017). Thus, *s_h_* represents the severity of CI, or the reduction in hatch rate due to CI in pairings between *Wolbachia*-negative ova and *Wolbachia*-positive sperm.

First, we consider *Wolbachia* strains that do not cause CI (*s_h_* = 0), which produce a stable equilibrium (*p*^) balanced by positive host fitness effects (*F* > 1) and imperfect maternal transmission (*µ* > 0) (Figure 1). When initially rare, *Wolbachia* must generate *F*(1 – *µ*) > 1 to spread deterministically from low frequency, regardless of whether they cause CI. Here, we assume *F*(1 – *µ*) > 1 given the spread and persistence of non-CI causing *Wolbachia* strains in nature (Hoffmann and Turelli 1997; Kriesner et al. 2013, 2016; Hamm et al. 2014; Kriesner and Hoffmann 2018; Meany et al. 2019). When *F*(1 – *µ*) < 1, rare infections will be deterministically lost (Hoffmann et al. 1990). If *F*(1 – *µ*) > 1, the stable equilibrium frequency is

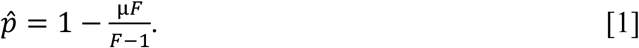

In this case, *p*^ increases from 0 to 1 – *µ* as *F* increases from 1/(1 – *µ*).

Next, we considered equilibria that incorporate CI (*s_h_* > 0) (Figure 1). Like non-CI strains, CI-causing strains must still increase host fitness to spread deterministically from low frequency such that *F*(1 – *µ*) > 1 (Turelli and Hoffmann 1991, 1995; Bakovic et al. 2018). CI-causing *Wolbachia* that generate *F*(1 – *µ*) > 1 and *Fµ* < 1 produce a single stable equilibrium between 0 and 1 given by

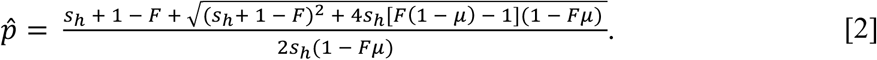

Assuming discrete host generations, the total population fraction of offspring in generation *t* + 1 can be divided with respect to whether mated males and females in generation *t* are carrying *Wolbachia*:

1. *Wolbachia*-*positive male and female*. In this case, there is a population of 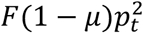 and also a population of 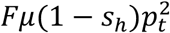 when ova become uninfected due to imperfect maternal transmission, resulting in a population fraction of 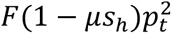 .
2. *Wolbachia*-*negative male and female*. The population fraction is simply (1 − *p_t_*)^2^.
3. *Wolbachia-negative male, Wolbachia-positive female*. Accounting for fecundity in *Wolbachia*-positive females, the population fraction is then *Fp_t_*(1 − *p_t_*).
4. *Wolbachia-positive male, Wolbachia-negative female*. The population from generation *t* is *p_t_*(1 − *p_t_*). A fraction 1 − *s_h_* of this population survives from this incompatible pairing, so the population fraction for generation *t* + 1 is *p_t_*(1 − *p_t_*)(1 − *s_h_*).

Assuming equal infection frequencies in males and females, the adult infection frequency in generation *t*, denoted *p_t_*, changes between generations as follows:

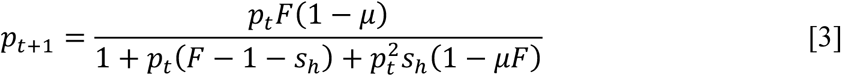

(Hoffmann et al. 1990). In the absence of CI (*s_h_* = 0), this model becomes

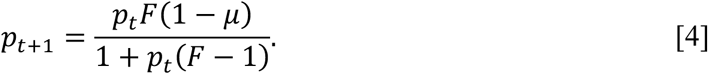

For a stability analysis of [4], see Appendix 1.

### Simulations to approximate temporal Wolbachia dynamics under finite population size

Following Turelli and Barton (2022) and Turelli et al. (2022), we approximated the stochasticity produced by finite population size using a stochastic transition matrix analogous to a haploid Wright-Fisher model of genetic drift, which assumes that generations are discrete and the new generation is a random sample of constant population size drawn from the total number of offspring produced by the previous generation (Crow and Kimura 1970). The model uses an effective population size of *N* females. The finite-population stochasticity, modeled as binomial sampling, is superimposed on the deterministic equilibria dynamics described by [3].

We now derive a model for modeling infection frequency *p*_0_ of *Wolbachia*-positive hosts which includes randomness effects from a finite female population of size *N*, *I*_0_ = *Np*_0_ of which are infected. We assume females produce the same number of eggs with each mating and that *N* does not change over generations. Starting with the current adult (female) infection frequency *p*_0_, the infection frequency among viable gametes in the next generation is determined by [3]. Then, the infection frequency in the next generation of *N* adult females is obtained from binomial sampling of this deterministic projection.

First, we compute the total number of *Wolbachia*-positive adult offspring in generation *t* + 1 in the absence of CI (*s_h_* = 0). For *Wolbachia*-positive females, we note that there are *FI*_0_ova produced. Each ovum has a probability 1 − *μ* of being infected, so the total number of infected offspring can be represented by a binomial random variable

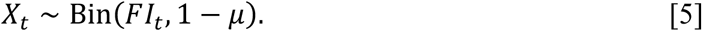

For the total number of offspring produced, we observe that all *Wolbachia*-negative females produce *Wolbachia*-negative ova. This gives a total of *N* − *I Wolbachia*-negative offspring for the next generation. For *Wolbachia*-positive females, we noted that *FI_t_* offspring are produced from matings. This gives a total population of *N* + (*F* − 1)*I_t_* offspring. The proportion of *Wolbachia*-positive adults for generation *t* + 1 is then

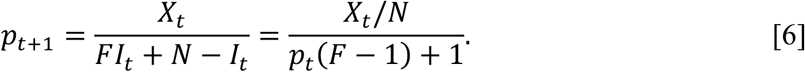

From [6], the expected value and variance of *p_t_*_+1_, given knowledge of *p_t_*, are readily computed as

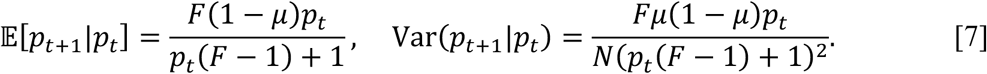

From the law of large numbers, as the population *N* → ∞, the variance in [7] tends to 0, and the random recursion [6] converges to the deterministic model [4]. See Appendix 2 for an analysis of long-term dynamics.

Now, allowing for CI (*s_h_* > 0), we can derive a random recursion for *p*_0_. For each of the stages of mating, ova production, and the generation of adult offspring, we record total populations in Table S1. The total *Wolbachia*-positive adult offspring resulting from *Wolbachia*- positive male and female matings is given by *X*_1_ ∼ Bin(*FpI*, 1 − *μ*). For *Wolbachia*-negative males and -positive females, total *Wolbachia*-positive offspring is *X*_2_ ∼ Bin(*F*(1 − *p*)*I*, 1 − *μ*). The total number of offspring is found by summing the four terms in the row labeled “adult offspring after CI” in Table S1. Denoting the normalized fractions *X̄_i_* = *X_i_*/*N*, the fraction of *Wolbachia*-positive offspring over the total population is then

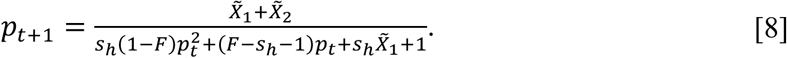

As *N* → ∞, from the law of large numbers [8] approaches the deterministic model [3].

Let us also consider the case where a subpopulation of “low transmitter” *Wolbachia*- positive females with high rates of imperfect maternal transmission are included in the host population. Here, the host population consists of *M* groups with group fractions (*r*_1_, …, *r_M_*). Rather than a single *μ* value, the *i*th subpopulation assumes a value *μ*_2_ for *i* = 1, …, *M*. To compute *Wolbachia*-positive offspring, we are now summing *M* separate binomial distributions. However, most computations from above carry over. For each group *i*, we compute *X_i_* ∼ Bin(*FIr_i_*, 1 − *μ_i_*). The number of *Wolbachia*-positive female eggs then becomes 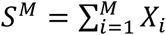 . For no CI, the total population is still *FI* + *N* − *I*, and

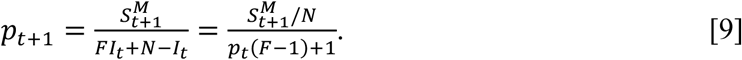

When *μ_i_* ≡ *μ* for all *i* = 1, …, *M*, then *S^M^* ∼ Bin(1 − *μ*, *FI*) and we recover [6]. Expected values are also straightforward to compute, with

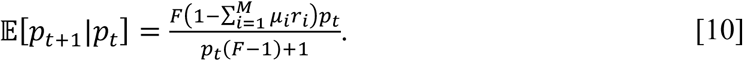

Each of the *Wolbachia*-positive populations *X_i_* for *i* …, *M* terms are independent, so by additivity of variance,

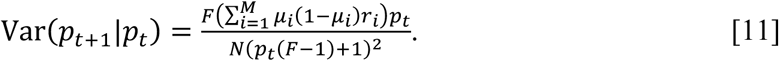

The stable equilibrium, where *p̂* = 𝔼[*p_t_*_+1_ |*p*_0_ = *p̂*], is very similar to [1]. Accounting for multiple *μ_i_* values, we have

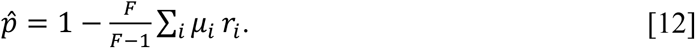

Finally, to generalize [8], we replace *X*_1_ and *X*_2_ with sums of independent binomials {*Y*_2_}_1≤*i*≤*M*_and {*Z_i_*}_1≤*i*≤*M*_, with

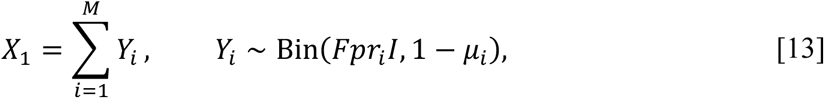

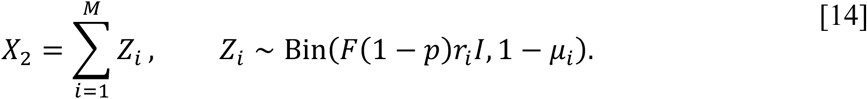

We also consider the case where host fitness effects fluctuate from one host generation to the next as independent, identically distributed log-normal random variables (Turelli et al. 2022). Here, we assume that each generation *F* is chosen independently from a log-normal distribution such that *F* = *e^X^*, where *X* is a normal random variable with mean μ_X_ and variance *σ*^2^ (Turelli et al. 2022). This implies that *F* has median *m* = *e^µ_x_^*^)^ and a squared coefficient of variation 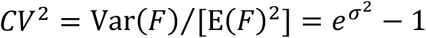. To produce a particular median, *m*, and *CV* for *F*, we set µ_X_ = ln(*m*) and *σ*^2^ = ln (*CV*^2^ + 1).

### Analysis of biologically plausible parameters values

To explore how *µ*, *F*, *s_h_*, and *N* values influence the size of temporal *Wolbachia* fluctuations, we implemented simulations across a broad range of plausible parameter values informed by empirical estimates for *w*Mel-like *Wolbachia* and field-collected *Drosophila* hosts. For imperfect maternal transmission, we explored a range of *µ* values based on maternal transmission estimates using field-collected females, as well as females reared under varying conditions in the lab. Estimates of *µ* (±BC_a_ confidence intervals) for *w*Mel from field-collected *D. melanogaster* range from low (*µ* = 0.026 [0.008, 0.059]; Hoffmann et al. 1998) up to moderate levels of imperfect transmission (*µ* = 0.11 [0.07, 0.17]; Olsen et al. 2001). Estimates generated in the lab range as high as *µ* = 0.415 (0.332, 0.493) when females are reared in the cold (20°C), as opposed to a standard temperature (25°C) (Hague et al. 2022, 2024). Estimates of *µ* for *w*San from field-collected *D. santomea* on São Tomé are moderate (*µ* = 0.068 [0.027, 0.154]), whereas *w*Yak in *D. yakuba* range from low at low altitude (*µ* = 0.038 [0.003, 0.184]) to moderate at high altitude (*µ* = 0.20 [0.087, 0.364]) (Hague et al. 2020*b*). Estimates of *µ* in the lab for *w*Yak range as high as *µ* = 0.15 (0.109 0.196) when females are reared in the cold (Hague et al. 2020*b*). Based this information, we explored a broad range of *µ* values from 0.001 ≤ *µ* ≤ 0.3.

Previous estimates of *µ* from a large sample of field-collected female *D. santomea* (*N* = 62) and *D. yakuba* (*N* = 71) on São Tomé in 2018 suggest that the majority of females have perfect or near perfect transmission rates, but a small proportion of “low transmitters” have high *µ* values. For *w*San, the total sample of *Wolbachia*-positive females had a mean value of *µ* = 0.068 (0.027, 0.154), but 8% of these females had a *µ* value greater than 0.5 with a mean of *µ* = 0.728. Similarly, for *w*Yak, the total sample of *Wolbachia*-positive females had a mean value of *µ* = 0.126 (0.062, 0.238), but approximately 11% of the females had a *µ* value greater than 0.5 with a mean of *µ* = 0.803 (Hague et al. 2020*b*). Other work using field-collected females found similar evidence of low transmitters for *w*Mel in *D. melanogaster* (Hoffmann et al. 1998), *w*Suz in *D. suzukii* (Hamm et al. 2014; see Figure 3), *w*Inn in *D. innubila* (Unckless et al. 2009), and *w*Ri in *D. simulans* (Carrington et al. 2011; see Figure 6). Hamm et al. (2014) termed females with high rates of imperfect *w*Suz transmission as “low transmitters” and we follow the same terminology here. Using the *D. yakuba*-clade data as a guide, we explored the effect of including a subpopulation of low transmitters (10% of *Wolbachia*-positive females) with a binomial *µ* value set to *µ* = 0.8. Here, the remaining 90% of *Wolbachia*-positive females have *µ* values set as described above (0.001 ≤ *µ* ≤ 0.3).

We know far less about *Wolbachia* effects on host fitness in natural populations due to the challenges of estimating *F* in the wild (Ross et al. 2019*b*). *Wolbachia* cells are found throughout host somatic tissue (Pietri et al. 2016), which is likely to have complex effects on components of host fitness in nature (Hoffmann et al. 1998; Weeks et al. 2007; Kriesner et al. 2016; Hague et al. 2020*a*, 2021; Cogni et al. 2021; Bruner-Montero and Jiggins 2023). We generally expect *F* > 1 based of the spread and persistence of *Wolbachia* strains in natural host populations (as described in the model above). Prior experiments found no evidence for fecundity effects of *w*Mel in *D. melanogaster* (Hoffmann et al. 1994) or *Wolbachia* in *D. yakuba*-clade hosts (Cooper et al. 2017); however, fecundity represents only one aspect of host fitness. Recent work suggests that *w*Mel blocks viruses in natural populations of *D. melanogaster* (Hedges et al. 2008; Teixeira et al. 2008; Cogni et al. 2021; Bruner-Montero and Jiggins 2023). *Wolbachia* effects on other measures of host fitness in the lab have ranged from positive (Brownlie et al. 2009; Russell et al. 2023) to negative (Hoffmann and Turelli 1988; Hoffmann et al. 1990; Kriesner et al. 2016) to context-dependent (Olsen et al. 2001; Fry et al. 2004). Beneficial *Wolbachia* effects of *F* > ∼1.2 (roughly a 20% relative fitness advantage for *Wolbachia*-positive females) have not been documented in any system (Weeks et al. 2007; Hamm et al. 2014; Meany et al. 2019; Hague et al. 2020*b*) and we generally expect values of *F* > 1.5 to be biologically unrealistic (Weeks et al. 2007; Hamm et al. 2014; Cooper et al. 2017; Meany et al. 2019; Hague et al. 2020*b*, 2022). Based on this information, we explored how *F* values ranging from 1 ≤ *F* ≤ 1.5 influence stochastic *Wolbachia* dynamics.

It is generally unknown whether host fitness effects fluctuate in natural host populations (Turelli et al. 2022); however, there is evidence that the fitness effects of *w*Mel on *D. melanogaster* fecundity (Kriesner et al. 2016) and virus-blocking (Chrostek et al. 2021) can depend on the environmental context. Therefore, we also ran simulations treating *F* as a log-normal random variable and explored scenarios with weakly (*CV* = 0.01) or strongly (*CV* = 0.1) fluctuating host fitness effects. To illustrate how these values correspond to fluctuating *F* values in our simulations, when median(*F*) = 1.05, a *CV* = 0.01 value corresponds to 2.5 and 97.5 percentiles of 1.03 and 1.071, respectively, and a *CV* = 0.1 value corresponds to 0.863 and 1.271, respectively.

We investigated three different values of *s_h_* based on estimates of CI strength from field-collected male *Drosophila*. Estimates using *D. melanogaster* males from central and northern Australia suggest that *w*Mel causes weak CI on the order of *s_h_* = 0.05 (Hoffmann et al. 1998; Kriesner et al. 2016). CI strength declines rapidly with male age (Shropshire et al. 2021) and Reynolds and Hoffmann (2002) found that one-day-old males derived from wild-collected larvae and pupae produce hatch rates of 0.39 in incompatible crosses, with large confidence intervals (0.15, 0.64). Given this information, Kriesner et al. (2016) conjectured that *s_h_* values of 0 to 0.1 are plausible for *w*Mel, but *s_h_* > 0.1 is unlikely. We are not aware of CI estimates for *D. yakuba*- clade *Wolbachia* using field-collected males; however, estimates in the lab indicate these strains reduce egg-to-adult viability in CI crosses by about 10-20% (*w*San: *s_h_* = 0.15 [0.12, 0.18]; *w*Yak: *s_h_* = 0.16 [0.13, 0.20]; *w*Tei: *s_h_* = 0.2 [0.17, 0.23]), although the presence and intensity of CI can vary depending on host and *Wolbachia* genotype (Cooper et al. 2017). Based on these data, we explored the effects of no CI (*s_h_* = 0) and weak CI (*s_h_* = 0.1) on stochastic *Wolbachia* dynamics. We also considered strong CI (*s_h_* = 0.45) characteristic of field-collected male *D. simulans* carrying *w*Ri (Turelli and Hoffmann 1995; Carrington et al. 2011).

For each unique combination of *µ*, *F*, and *s_h_* values, we used the *symbiontmodeler* package to run a set of 25 replicate simulations. Each individual simulation began with an arbitrary intermediate *Wolbachia* frequency of *p*_0_ = 0.4 and ran for 10,000 host generations (similar *p*_0_ values do not qualitatively alter the results). We removed an initial burn-in of 500 host generations as *Wolbachia* spread from *p*_0_ = 0.4 to a stable equilibrium, and then calculated the mean (*p̅*) and standard deviation (*p*_SD_) of *Wolbachia* frequencies from generations 501- 10,000 for each simulation. We also ran each set of simulations with different plausible values for host population size (*N*) ranging from 10^3^ to 10^6^ (McKenzie 1980; McInnis et al. 1982; Powell 1997; Gravot et al. 2004; Bergland et al. 2014). Estimates of local population sizes of *Drosophila* made from mark-release-recapture methods report local census sizes on the order of 10^4^ to 10^5^ with considerable variation across time and location (McKenzie 1980; McInnis et al. 1982; Powell 1997; Gravot et al. 2004; Bergland et al. 2014).

Finally, to quantify the influence of the different parameter values on temporal *Wolbachia* dynamics, we modeled the *p̅* and *p*_SD_ values with linear effects for *N*, *µ*, the presence/absence of low transmitters, *F*, the presence/absence of fluctuating *F* values (i.e., treating *F* as fixed or a random variable), and *s_h_*. Because the infection frequency is always between zero and one, we modeled the distribution of the *p̅* and *p*_SD_ values as beta random variables. Models were implemented in R version 4.4.2 using the *mgcv* package version 1.9.1 (Wood 2017). We found that a small handful of simulations had infections that persisted at *p* > 0 for the full 10,000 host generations for combinations of parameter values that do not predict a stable equilibrium of *p̂* > 0 (e.g., *F* = 1.05, *µ* = 0.05, *s_h_* = 0). We removed these simulations from our regression analysis, because they are predicted to be deterministically lost over time and we did not want the *p̅* and *p*_SD_ values to bias our analyses of stable equilibria. Finally, we used the “plot_predictions” function from the *marginaleffects* package version 0.25 (Arel-Bundock et al. 2024) to visualize the results by generating model predictions for *p̅* and *p*_SD_ as a function of the *µ* value modulated by the presence/absence of low transmitters, *F*, and *s_h_*.

## RESULTS AND DISCUSSION

We used *symbiontmodeler* to run a total of 491,400 simulations across 3,276 unique parameter combinations informed by empirical estimates from *Wolbachia*-*Drosophila* systems and summarized the mean (*p̅*; Figure 2) and standard deviation (*p*_SD_; Figure 3) of *Wolbachia* frequencies from each individual simulation. We then used multiple regression to summarize how different values of host population size (*N*), imperfect maternal transmission (*µ*), host fitness effects (*F*), and CI strength (*s_h_*) influence *Wolbachia* frequencies (measured as *p̅*) and the size of stochastic *Wolbachia* fluctuations over time (measured as *p*_SD_) in simulated host populations (Figures 4, 5, Tables S2, S3). We first briefly detail how the mean *Wolbachia* frequencies from our stochastic simulations align with equilibria predictions from the strictly deterministic model (Figure 1). Then, we examine the influence of different model parameters on the size of stochastic *Wolbachia* fluctuations, as well as how other factors might contribute to temporal fluctuations in natural host populations.

**Figure 2.**
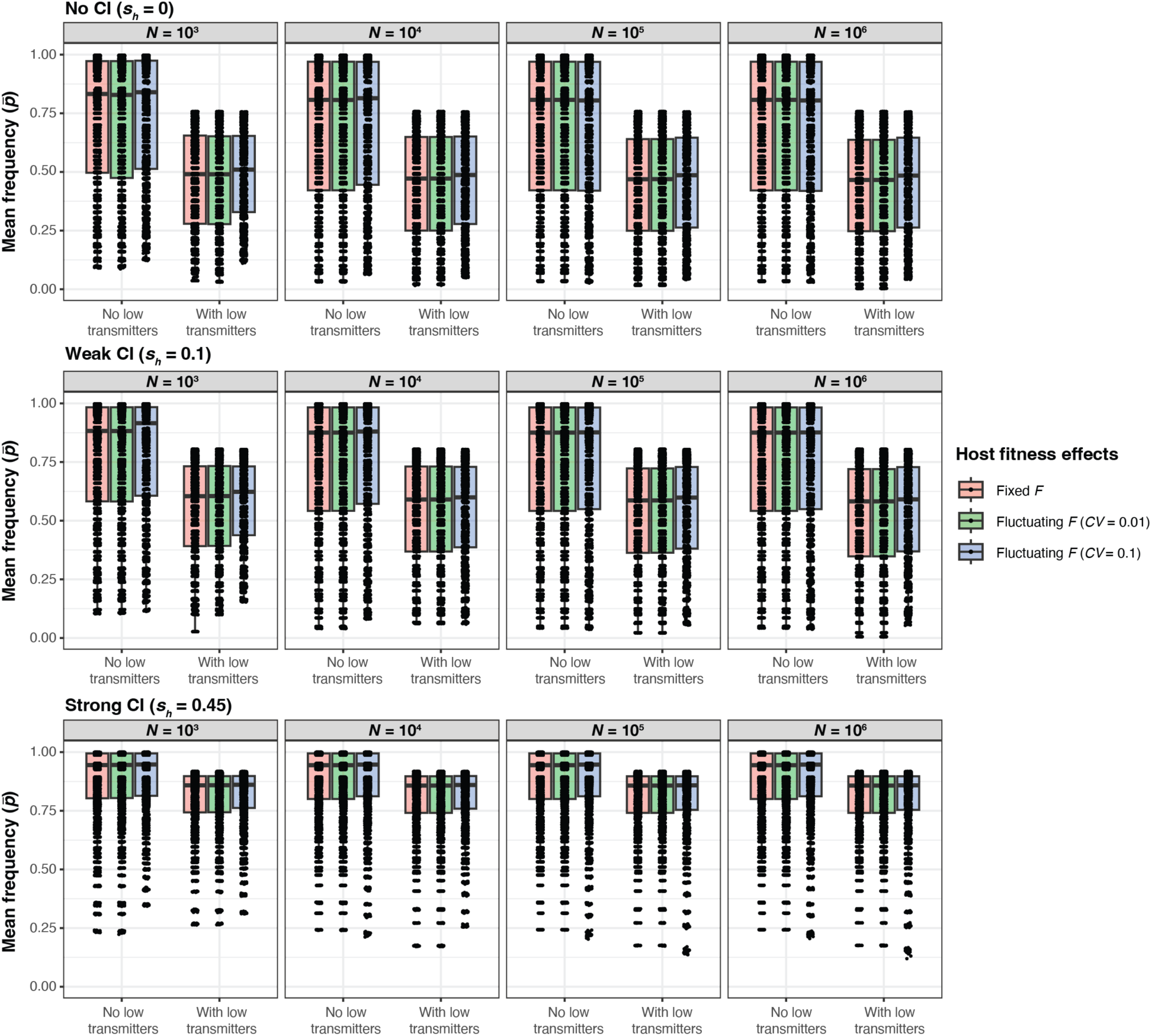
Results from individual simulations exploring the contributions of *N*, *µ*, *F*, and *s_h_* to mean *Wolbachia* frequencies (*p̅*). Each boxplot represents the distribution of *p̅* values from simulations implemented with different combinations of *µ* (0.001 ≤ *µ* ≤ 0.3) and *F* (1 ≤ *F* ≤ 1.5). Datasets are further separated based on whether low transmitters and fluctuating host effects (*CV* = 0.01 or 0.1) are present. Results are shown for different CI strengths (none, weak, or strong) and population sizes (10^3^, 10^4^, 10^5^, 10^6^). Each point represents a *p̅* value from an individual simulation. The first 500 generations were removed from each simulation before calculating *p̅*.

**Figure 3.**
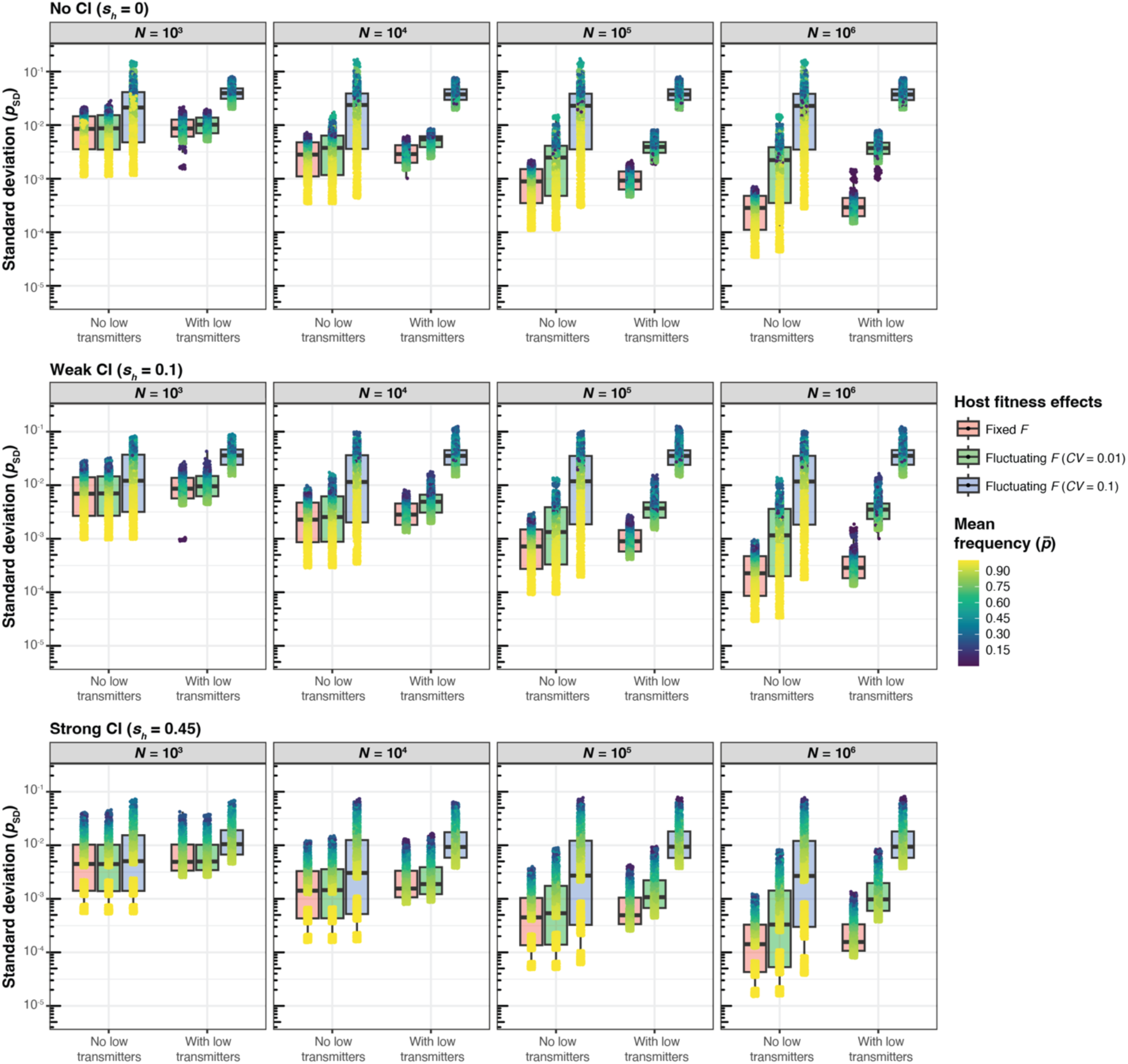
Results from individual simulations exploring the contributions of *N*, *µ*, *F*, and *s_h_* to the standard deviation of *Wolbachia* frequencies (*p*_SD_). Each boxplot represents the distribution of *p*_SD_ values from simulations implemented with different combinations of *µ* (0.001 ≤ *µ* ≤ 0.3) and *F* (1 ≤ *F* ≤ 1.5). Datasets are further separated based on whether low transmitters and fluctuating host effects (*CV* = 0.01 or 0.1) are present. Results are shown for different CI strengths (none, weak, or strong) and population sizes (10^3^, 10^4^, 10^5^, 10^6^). Each point represents a *p*_SD_ value from an individual simulation and is color-coded to indicate mean *Wolbachia* frequency (*p̅*) from the simulation. The first 500 generations were removed from each simulation before calculating *p*_SD_.

**Figure 4.**
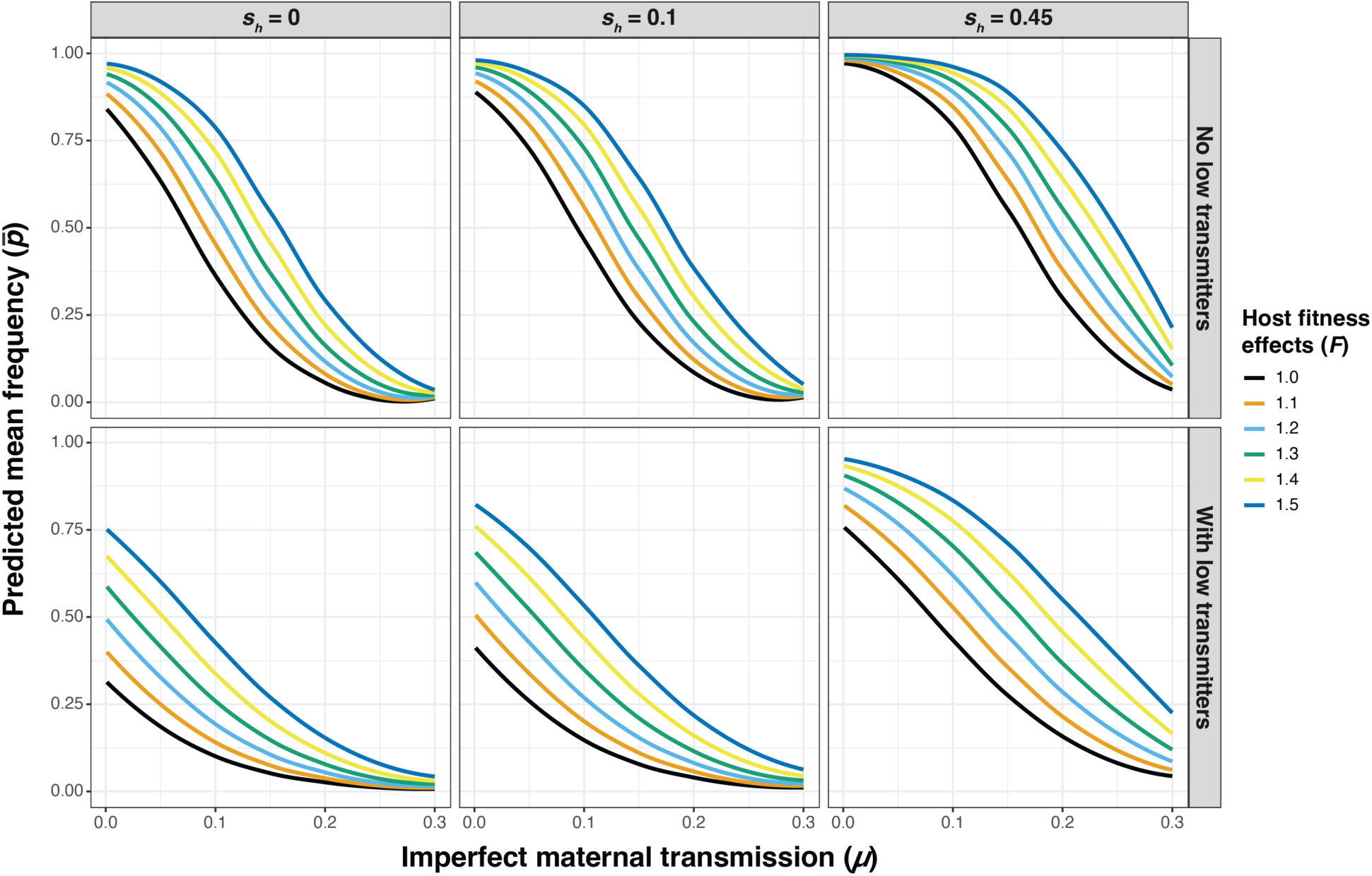
Predicted mean *Wolbachia* frequency (*p̅*) values from the regression analysis as a function of imperfect maternal transmission (*µ*) and conditioned by the presence/absence of low transmitters, strength of host fitness effects (*F*), and cytoplasmic incompatibility (*s_h_*). Note that prediction lines average the effects of host population size (*N*) and fixed/fluctuating host effects (*CV* = 0.01 and 0.1).

**Figure 5.**
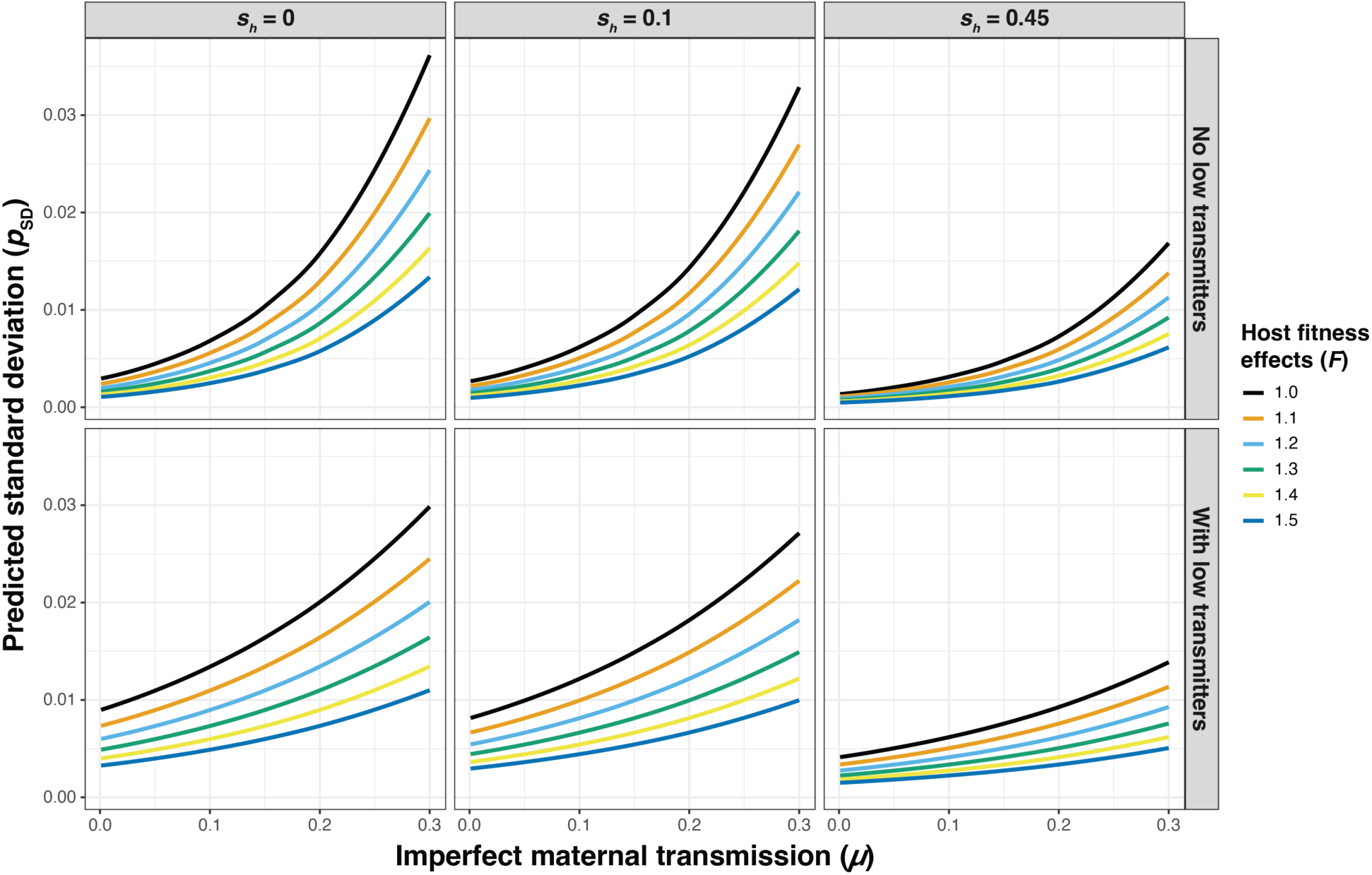
Predicted standard deviation (*p*_SD_) values of *Wolbachia* frequencies over time from the regression analysis as a function of imperfect maternal transmission (*µ*) and conditioned by the presence/absence of low transmitters, strength of host fitness effects (*F*), and cytoplasmic incompatibility (*s_h_*). Note that prediction lines average the effects of host population size (*N*) and fixed/fluctuating host effects (*CV* = 0.01 and 0.1).

Expectedly, the *p̅* values from our simulations (Figures 2, 4) closely align with equilibria predictions (*p̂*) from the deterministic model shown in equation [2] (Figure 1), which demonstrates that high rates of maternal transmission (i.e., smaller *µ* values), strong beneficial host effects, and strong CI yield high equilibrium frequencies. Here, we present simulation results from a population size of *N* = 10^4^ as a simple case to illustrate how mean *Wolbachia* frequencies are determined by parameter values of *µ*, *F*, and *s_h_* (Figure 6A). Within the range of plausible *µ*, *F*, and *s_h_* values we explored, the regression analysis indicated that the value of *μ* had the strongest relative effect on *p̅* (*β* = −22.4, *P* < 0.001), such that increasing rates of imperfect maternal transmission decrease *Wolbachia* frequencies (Table S2). The value of *F* had a smaller relative effect than *μ* on *p̅* (*β* = 3.77, *P* < 0.001), as strong host fitness benefits increase *Wolbachia* frequencies. Finally, the addition of weak CI (*s_h_* = 0.1) had a small effect on *p̅* (*β* = 0.427, *P* < 0.001), whereas strong CI characteristic of strains like *w*Ri (*sh* = 0.45) had a stronger effect (*β* = 1.92, *P* < 0.001) as the frequency dependent benefit of CI drives *Wolbachia* to high frequencies.

**Figure 6.**
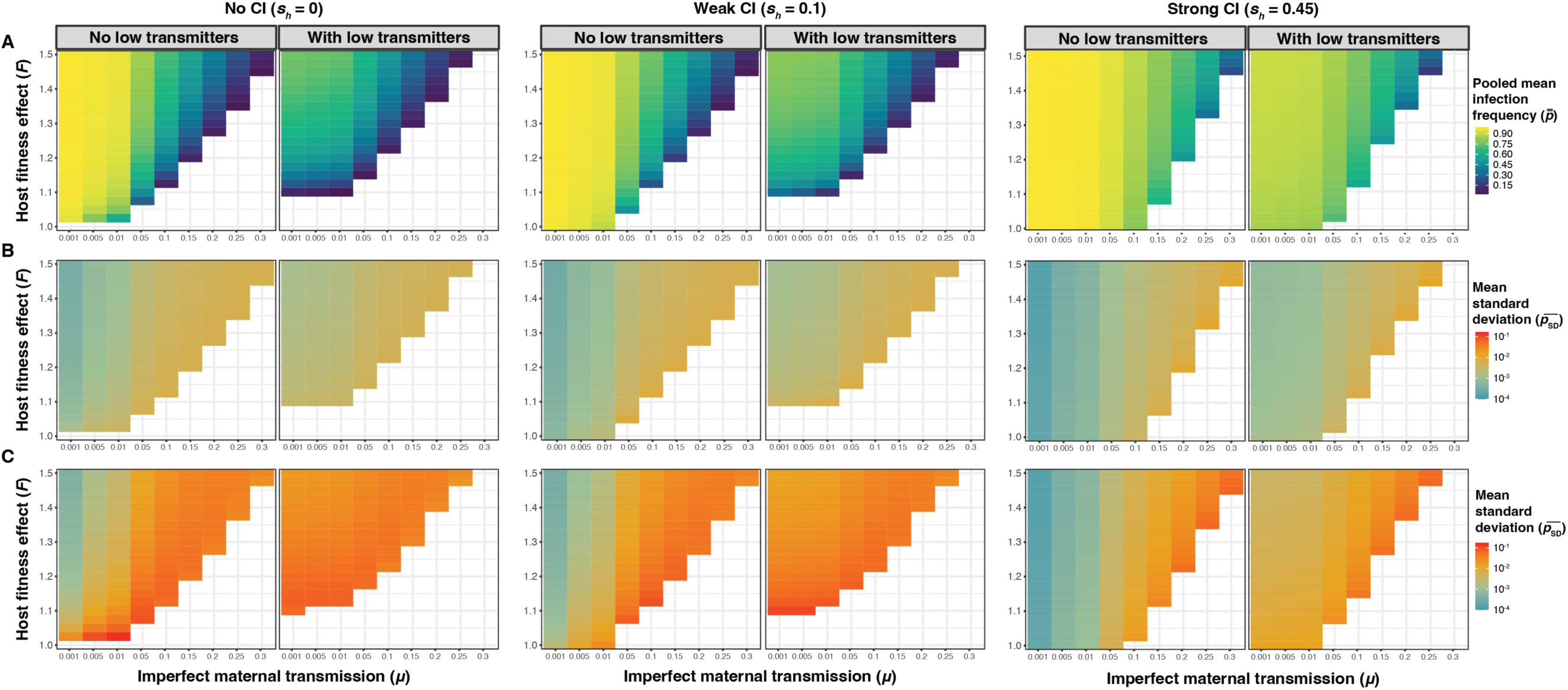
Summary of results from simulations exploring the contributions of *µ*, *F*, and *s_h_* to the mean (*p̅*) and standard deviation (*p*_SD_) of *Wolbachia* frequencies from simulations with a finite host population size of *N* = 10^4^. (A) Each cell represents the pooled mean *Wolbachia* frequency (*p̿*) from 25 replicate simulations with a set *µ* and *F* value. **(B)** Each cell represents the mean standard deviation 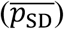 of *Wolbachia* frequencies from 25 simulations with a set *µ* and *F* value. Note that 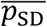 values are plotted on a log scale. **(B)** Each cell represents 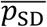 from 25 simulations with a set *µ* and *F* value with strongly fluctuating fitness effects (*CV* = 0.1). Datasets are further separated based on the presence/absence of low transmitters and CI strength (none, weak, or strong).

The inclusion of a subpopulation of low transmitter females (10% of *Wolbachia*-positive females with *µ* = 0.8) was also associated with lower *p̅* values (*β* = −2.43, *P* < 0.001). We also found an interaction effect between the presence of low transmitters and the *μ* value (*β* = 8.26, *P* < 0.001), such that the rate of decrease in *p̅* as a function of *μ* is slightly less when low transmitters are present (Figure 4). While including a subpopulation of low transmitter females generally reduced mean *Wolbachia* frequencies, we note that this effect was strongest for *Wolbachia* strains that do not cause CI (e.g., *w*Mel in *D. melanogaster*). In the *N* = 10^4^ example (Figure 6), the no-CI simulations (*s_h_* = 0) without low transmitters had a median *p̅* value of 0.807, which dropped down to 0.472 in the simulations with low transmitters. In contrast, the median *p̅* value dropped from 0.875 without low transmitters to 0.591 with low transmitters for weak CI (*s_h_* = 0.1) and from 0.945 to 0.858 for strong CI (*s_h_* = 0.45). The impact of low transmitters on *p̅* is especially strong for relatively small values of *µ* (≤ 0.01) and *F* (≤ 1.15), which are otherwise predicted to generate high equilibrium frequencies (Figure 6). For instance, the values of *µ* = 0.001, *F* = 1.1, and *s_h_* = 0 generated a pooled mean of *p̿* = 0.989 across 25 replicate simulations, but the addition of low transmitters dropped the mean down to *p̿* = 0.11.

Studies using field-collected females show that low transmitters are present in some host populations (Hoffmann et al. 1998; Unckless et al. 2009; Carrington et al. 2011; Hamm et al. 2014; Hague et al. 2020*b*), but it is generally unclear why heterogeneity in maternal transmission rates exists among individual hosts. Low transmitters could develop if a subpopulation of hosts experience extreme environmental conditions (e.g., a cold bout) that reduce *Wolbachia* densities and perturb maternal transmission (Ulrich et al. 2016; Ross et al. 2017, 2019*a*; Foo et al. 2019; Hague et al. 2020*b*, 2020*a*, 2022, 2024; Chrostek et al. 2021). Thermal conditions can vary on small spatial scales (e.g., individual fruits), so it is plausible that individual females within a population could experience different environmental conditions that impact maternal transmission (Roberts and Feder 2000; Potter et al. 2009; Saudreau et al. 2009; Pincebourde and Woods 2012; Woods et al. 2015). Heterogeneity in *Wolbachia* densities and/or transmission could also arise due to variation in host diet (Serbus et al. 2015; Christensen et al. 2019), age (Reynolds and Hoffmann 2002; Shropshire et al. 2021), and the host and *Wolbachia* genomes (Gu et al. 2022; Hague et al. 2022). For instance, densities of the *w*Inn strain in *D. innubila* host individuals varied by 20,000 fold and correlated with maternal transmission rates in populations sampled in Arizona (Unckless et al. 2009).

Next, we used the regression analysis to summarize how different parameter values contribute to the size of stochastic *Wolbachia* fluctuations over time, measured as *p*_SD_ (Figure 5, Table S3). Again, the value of *μ* had the strongest relative effect on increasing *p*_SD_ (*β* = 8.51, *P* < 0.001), such that higher rates of imperfect maternal transmission tend to generate larger *Wolbachia* fluctuations. To a lesser extent, the inclusion of low transmitters also increased *p*_SD_ values (*β* = 1.12, *P* < 0.001). We also found an interaction effect between the value of *μ* and the presence of low transmitters (*β* = −4.42, *P* < 0.001), where the rate of increase in *p*_SD_ as a function of *μ* is smaller when low transmitters are present. Beyond maternal transmission rates, the results illustrate an intuitive pattern where strong selection for *Wolbachia* arising from *F* and *s_h_* is associated with higher *Wolbachia* frequencies (*p̅*) and smaller stochastic fluctuations (*p*_SD_) (Figure S1). The value of *F* had a smaller relative effect than *μ* on *p*_SD_ (*β* = −2.04, *P* < 0.001). If we assume fluctuating host effects and treat *F* as a log-normal random variable, weak *F* fluctuations (*CV* = 0.01) only have a minor effect on increasing *p*_SD_ values (*β* = 0.423, *P* < 0.001), whereas strong fluctuations predictably have a larger effect (*β* = 1.78, *P* < 0.001). The addition of weak CI (*s_h_* = 0.1) had a relatively small effect on decreasing *p*_SD_ values (*β* = −9.85 x 10^-2^, *P* < 0.001). Stronger CI (*s_h_* = 0.45) had a larger effect on *p*_SD_ (*β* = −0.783, *P* < 0.001), as strong CI is associated with high, stable infection frequencies. Host population size (*N*) had only a small relative effect (*β* = −4.14 x 10^-7^, *P* < 0.001), although *p*_SD_ predictably declined as population size increased.

Broadly, our simulation results are consistent with empirical observations of temporal *Wolbachia* dynamics in natural host populations. *w*Mel-like strains like *w*Mel and *w*San that show evidence of imperfect maternal transmission and cause weak or no CI also tend to fluctuate temporally and spatially in host populations (Hoffmann et al. 1994; Kriesner et al. 2016; Cooper et al. 2017; Hague et al. 2020*b*). For instance, estimates of *w*Mel maternal transmission rates from field-collected female *D. melanogaster* have been recorded as high as *µ* = 0.11 (0.07, 0.17) and higher values can be generated in the lab by rearing hosts in the cold (Olsen et al. 2001; Hague et al. 2022, 2024). Estimates of CI strength for *w*Mel from field-collected males crossed with *Wolbachia*-negative laboratory females are consistent with *s_h_* ≈ 0 (unless males are very young) (Hoffmann et al. 1998; Reynolds and Hoffmann 2002; Kriesner et al. 2016; Shropshire et al. 2021). *w*Mel frequencies have been shown to fluctuate significantly over time in host populations in Australia (Hoffmann et al. 1998) and across geography in Australia, North America, and Africa (Kriesner et al. 2016). Similar levels of imperfect maternal transmission have been recorded for *w*San from field-collected *D. santomea* (*µ* = 0.120, [0.032, 0.346]) and for *w*Yak from *D. yakuba* (*µ* = 0.20 [0.087, 0.364]) on São Tomé (Hague et al. 2020*b*). *w*San and *w*Yak cause weak CI in the lab (*s_h_* = 0.15 [0.12, 0.18] and *s_h_* = 0.16 [0.13, 0.20], respectively) and their frequencies have both fluctuated temporally on the island of São Tome (Cooper et al. 2017; Hague et al. 2020*b*). *w*Yak frequencies also vary by altitude on São Tome (Hague et al. 2020*b*). These patterns are by no means characteristic of all *w*Mel-like *Wolbachia*, as some *w*Mel-like strains do cause strong CI in their native hosts (e.g., *w*Seg in *D. seguyi*; Shropshire et al. 2024). Other, more diverged *Wolbachia*, like the *w*Ri-like *w*Suz strain in *D. suzukii* are imperfectly transmitted (*µ* = 0.14 [0.04, 0.27]), do not cause CI, and persist at intermediate frequencies that vary across time and space (Hamm et al. 2014; Turelli et al. 2018).

Notably, across the range of parameter values we explored, declining rates of maternal transmission (i.e., increasing values of *µ*) had the strongest relative effect on increasing the size of temporal *Wolbachia* fluctuations, as compared to the strength of host fitness effects and CI (Table S3). These results are interesting in light of the fact that maternal transmission rates of the *w*Mel-like strains *w*Mel and *w*Yak are temperature dependent. Both strains exhibit relatively high rates of transmission when hosts are reared in the lab at a standard 25°C (*w*Mel: *µ* = 0.056 [0.022, 0.122], *w*Yak: *µ* = 0.010 [0, 0.034]), but transmission rates decline when hosts are reared in the cold at 20°C (*w*Mel: *µ* = 0.197 [0.130, 0.269], *w*Yak: *µ* = 0.150 [0.106, 0.196]) (Hague et al. 2020*b*, 2024). These results suggest the stochastic fluctuations of *w*Mel-like *Wolbachia* may increase in size under conditions where the maternal transmission rate is perturbed by the environment.

The fluctuating patterns of non-CI-causing strains like *w*Mel contrast those of strong-CI-causing *Wolbachia* like *w*Ri in *D. simulans*, which persists at high, stable infection frequencies in nature. *w*Ri generates strong CI (*s_h_* ≈ 0.45) and seems to have relatively high rates of maternal transmission in the field (e.g., *µ* = 0.026 [0.004, 0.057]) and perfect transmission in the lab across different temperatures (Hoffmann et al. 1990; Turelli and Hoffmann 1995; Carrington et al. 2011; Kriesner et al. 2013; Hague et al. 2024). Similarly, the CI-causing strains *w*Ha (in *D. simulans*) and *w*Ana (in *D. ananassae*) persist at high, stable infection frequencies (Rousset and Solignac 1995; Bourtzis et al. 1996; James and Ballard 2000; Ballard 2004; Choi et al. 2015) and have thermally stable maternal transmission rates in the lab (Hague et al. 2024).

Given that some instances of large temporal fluctuations of *w*Mel-like *Wolbachia* have been observed (e.g., >0.7 change in frequency), we also examined the specific combinations of parameters that generate the largest stochastic fluctuations in our simulations. Here, the specific parameters that generated the largest *p*_SD_ values do not strictly align with the general trends outlined above (Figure 5). Again, we focus on the population size of *N* = 10^4^ as a simple case (Figure 6). Assuming strong fluctuating effects on host fitness (*CV* = 0.1), the simulations with a relatively low rate of imperfect maternal transmission (*μ* = 0.01, no low transmitters), weak host benefits (*F* = 1.025), and no CI (*s_h_* = 0) generated the highest average *p*_SD_ value of 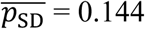 across 25 replicate simulations (*p̿* = 0.540) (Figure S2). In this region of parameter space, no CI and the presence of strongly fluctuating *F* values with a median slightly >1 cause *Wolbachia* to fluctuate between strongly favored (*F*[1 – *µ*] > 1) and disfavored (*F*[1 – *µ*] < 1) in the host population (Figure S2). A median of value of *F* = 1.025 with *CV* = 0.1 generates 2.5 and 97.5 percentiles of 0.843 and 1.251, respectively. These results highlight how a unique set of conditions—particularly fluctuating host effects with a median *F* value near one—are required to generate the largest stochastic fluctuations in our model.

Do host fitness effects fluctuate in natural host populations? Little is known about *Wolbachia* effects on host fitness in nature, although the endosymbionts are generally considered to be beneficial based on the spread and persistence of *Wolbachia* in natural host populations (Weeks et al. 2007; Hamm et al. 2014; Cooper et al. 2017; Meany et al. 2019; Ross et al. 2019*b*; Hague et al. 2020*b*, 2022; Hoffmann and Cooper 2024). While we do not know if host fitness effects fluctuate (Turelli et al. 2022), there is evidence that the effects of *w*Mel on *D. melanogaster* fecundity (Kriesner et al. 2016) and virus-blocking (Chrostek et al. 2021) can depend on the environmental context. The non-CI-causing strain *w*Au in *D. simulans* has also been shown to have context-dependent host effects depending on where host breeding occurs (Cao et al. 2019). Given that temperature has pervasive effects on *Wolbachia* densities in host tissues (Hague et al. 2020*a*, 2024), it is also plausible that temperature-induced changes to *Wolbachia* densities could in turn alter *Wolbachia* effects on various host fitness components.

Our results here serve as a baseline for understanding the contribution of finite-population stochasticity to temporal *Wolbachia* dynamics for *w*Mel-like *Wolbachia*. The dynamics we modeled here may provide a plausible explanation for examples of minor frequency fluctuations, such as temporal variation of *w*Mel in temperate Australia in 1993-96 (Figure S3) (Hoffmann et al. 1998) or fine-scale geographic variation of *w*Yak on the island of São Tomé in 2018 (Figure S4) (Hague et al. 2020*b*). However, the additional assumption of fluctuating host fitness effects is required to explain examples of larger temporal fluctuations (e.g., >0.4 change in frequency), such as *w*Mel variation in subtropical Australia in 1995-96 (Figure S5) (Hoffmann et al. 1998) or *w*San variation on São Tomé in 2005-18 (Figure S5) (Hague et al. 2020*b*).

It is likely that other factors contribute to temporal fluctuations of *w*Mel-like *Wolbachia*, especially given that large fluctuations (e.g., >0.6 change in frequency) have been observed over sampling periods as short as a month (e.g., Figure S5) (Hoffmann et al. 1998). What else could cause *w*Mel-like *Wolbachia* frequencies to fluctuate over time? Perturbation of the maternal transmission rate, for example, due to a change in environmental conditions over time, could cause the equilibrium frequency to decline (Hague et al. 2020*b*, 2022, 2024). In a similar fashion, temporal changes to the strength of host fitness effects or CI could alter *Wolbachia* equilibria, as these parameter can also very depending on environmental conditions, host age, and/or the *Wolbachia* and host genomes (Hoffmann et al. 1990; Clancy and Hoffmann 1998; Olsen et al. 2001; Reynolds and Hoffmann 2002; Bordenstein and Bordenstein 2011; Versace et al. 2014; Kriesner et al. 2016; Cooper et al. 2017; Cao et al. 2019; Shropshire et al. 2021). The model also assumes that hosts have discrete generations and a single, fixed population size. Aspects of insect population dynamics and ecology, such as seasonal variation in population size, could contribute to fluctuating *Wolbachia* frequencies over time (Rasgon and Scott 2004; Farkas and Hinow 2010; Turelli 2010; Hancock et al. 2011; Turelli and Barton 2022). Host migration among structured populations with different *Wolbachia* equilibria dynamics could also plausibly alter frequencies within a given focal population (Hoffmann et al. 1998; Flor et al. 2007; Jansen et al. 2008; Engelstädter and Telschow 2009; Haygood and Turelli 2009; Hancock et al. 2011).

In summmary, our simulations provide a baseline for understanding the stochasticity of temporal *Wolbachia* dynamics, highlighting how non-CI-causing *w*Mel-like strains with relatively poor maternal transmission rates are more likely to fluctuate over time than strong CI-causing strains like *w*Ri. These results motivate further work to explore the incidence and causes of endosymbiont frequency fluctuations in natural host populations. The temporal dynamics of *w*Mel are quite variable at certain locations, but on a broader geographic scale, *w*Mel frequencies persist along a stable latitudinal cline in eastern Australia, raising further questions about the temporal and geographic scales at which *Wolbachia* frequencies vary (Hoffmann et al. 1994, 1998; Kriesner et al. 2016). Studies of *Wolbachia* dynamics within individual host populations on a fine temporal or geographic scale are still relatively uncommon (Turelli and Hoffmann 1995; Hoffmann et al. 1998; Hamm et al. 2014; Kriesner et al. 2016; Cooper et al. 2017; Hague et al. 2020*b*; Wheeler et al. 2021). Moving forward, the publicly available R package *symbiontmodeler* can be used to model how field estimates of maternal transmission, fecundity, and CI at different timepoints or locations contribute to variation in the prevalence of *Wolbachia*, as well as other symbionts (e.g., *Cardinium*; Perlman et al. 2008; Harris et al. 2010). Uncovering sources of temporal and spatial variation in *Wolbachia* prevalence is ultimately critical for predicting endosymbiont spread and the long-term evolutionary dynamics of endosymbiotic relationships.

## DATA ACCESSIBILITY STATEMENT

The *symbiontmodeler* R package and scripts to reproduce the regression analysis and figures will be made publicly available upon publication.

## AUTHOR CONTRIBUTIONS

JMG: funding acquisition (equal), methodology (equal), formal analysis (lead), software (lead), visualization (lead), writing – original draft (supporting), writing – review and editing (supporting). JK: methodology (equal), formal analysis (supporting), writing – original draft (supporting), writing – review and editing (supporting). MTJH: funding acquisition (equal), conceptualization (lead), supervision (lead), writing – original draft (lead), writing – review and editing (lead).

## ACKNOWLEDGEMENTS

This work was supported by the University of Scranton through a Faculty Internal Research Award to JMG and MTJH and a startup package awarded to MTJH from the College of Arts and Sciences Dean’s Office.

