## Supplemental Information for "Stochastic fluctuations of the facultative endosymbiont *Wolbachia* due to finite host population size"

### Appendix 1

For a stability analysis of [4], we define

$$g(x) = \frac{F(1-\mu)x}{x(F-1)+1}. \qquad \text{[A.1]}$$

Then $p_{t+1} = g(p_t)$, and stable equilibria occur for values $\hat{p}$ which satisfy $\hat{p} = g(\hat{p})$, which are found to occur at 0 and

$$\hat{p} = 1 - \frac{F\mu}{F-1}. \qquad \text{[A.2]}$$

In order for $\hat{p}$ to be contained in $(0,1)$ we require that $F > 1$, $\mu > 0$ and $F(1-\mu) > 1$. Since

$$g'(x) = F(1-\mu)\frac{1}{\left((F-1)x+1\right)^2}, \qquad \text{[A.3]}$$

$\hat{p}$ is a stable fixed point when

$$g'(\hat{p}) < 1 \quad \Rightarrow \quad F(1-\mu) > 1. \qquad \text{[A.4]}$$

We can strengthen this statement to show when $\hat{p}$ is a global attractor.

**Theorem**: If $F(1-\mu) > 1$, $\mu > 0$ and $F > 1$, then all iterations of the deterministic model [4] with initial conditions $p_0 \in (0,1]$ converge to the single fixed point $\hat{p}$.

*Proof.* We use the following properties of $g(x)$:

1. $g(0) = 0$,
2. $g'(0) = F(1-\mu) > 1$,
3. $g'(x) > 0$, or $g$ is increasing on $[0,1]$,
4. $g''(x) < 0$, or $g$ is concave in $[0,1]$.

This implies that in the interval $(0,\hat{p})$, $g(x)$ lies above the line $y = x$ and for $x \in (0,\hat{p})$, then $g(x) \in (x,\hat{p})$. In other words, the sequence of iterates $p_k = g(p_{k-1})$ with $p_0 \in (0,\hat{p})$ is an increasing sequence in $[p_0,\hat{p})$, and this must converge to $\hat{p}$ as $k \to \infty$.

For values in $(\hat{p},1]$, we note that since $g'(\hat{p}) < 1$ and $g'$ is decreasing, $g(x)$ must lie below the line $y = x$. Iterates beginning at $p_0 \in (\hat{p},1]$ are thus contained in $(\hat{p},1]$ and are decreasing. Again, this implies that they must converge to $\hat{p}$.

### Appendix 2

For analyzing the fluctuations of $p_t$ over many generations, we now approximate the behavior of the discrete recursion $\{p_t\}_{t\in\mathbb{N}}$ in [6] with an Ornstein-Uhlenbeck (OU) process, a continuous stochastic process $\{P_t\}_{t\geq 0}$ for mean-reverting behavior. For our problem, the stochastic differential equation for the OU process takes the form

$$dP_t = \kappa(\hat{p} - P_t)dt + \sigma dW(t). \tag{A.5}$$

In the drift term, $P_t$ converges to $\hat{p}$ at an exponential rate of $\kappa > 0$. For the noise term, $W(t)$ is a standard Brownian motion, with a volatility $\sigma > 0$. The benefit of estimating $p_t$ with [A.5] is that OU processes have exact expressions for the variance of a process after time $t$. Specifically, for a process starting at the deterministic equilibrium point $\hat{p}$, $\mathrm{Var}(P_t) = \frac{\sigma^2}{2\kappa}(1 - e^{-2\kappa t})$.

In our case, we already computed the fixed point $\hat{p} = 1 - \frac{F\mu}{F-1}$, and we will estimate $\kappa$ and $\sigma$ with equation [7]. This is done by discretizing time in [A.5], with a time step $\Delta t$ equal to a typical intergenerational period. One *Drosophila* generation corresponds to roughly one month in the field. The drift intensity can be approximated by the infinitesimal generator of $p_t$, given by

$$R(p) = \frac{\mathbb{E}[p_{t+1}|p_t = p] - p}{\Delta t} = \frac{1}{\Delta t}\left(\frac{F(1-\mu)p}{p(F-1)+1} - p\right). \tag{A.6}$$

If $p_t = \hat{p}$, then $\mathbb{E}[p_{t+1}|p_t] = \hat{p}$ and so $R(\hat{p}) = 0$. The first-order approximation of $R(p)$ near $\hat{p}$ is then

$$R(p) \approx R'(\hat{p})p \quad \Rightarrow \quad \kappa \approx -R'(\hat{p}). \tag{A.7}$$

From [7] and [1], we obtain the simplification

$$\frac{F(1-\mu)}{\hat{p}(F-1)+1} = 1. \tag{A.8}$$

This is used to compute

$$R'(\hat{p})\Delta t = \frac{F(1-\mu)}{\left((F-1)\hat{p}+1\right)^2} - 1 = \frac{1}{F(1-\mu)} - 1. \tag{A.9}$$

We thus obtain

$$\kappa \approx \frac{1}{\Delta t}\left(1 - \frac{1}{F(1-\mu)}\right). \tag{A.10}$$

Now for estimating $\sigma$, we write

$$\sigma^2 \Delta t \approx \mathrm{Var}(p_{t+1}|p_t = \hat{p}) = \frac{F\mu(1-\mu)\hat{p}}{N(\hat{p}(F-1)+1)^2} = \frac{\mu\hat{p}}{FN(1-\mu)}. \tag{A.11}$$

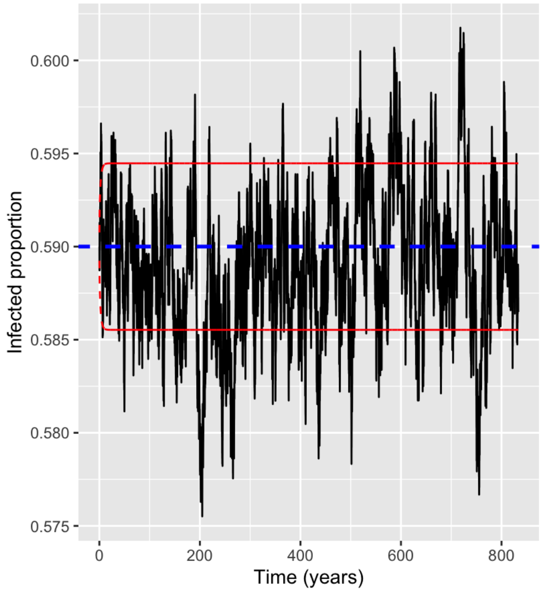


**Figure A.1** A simulation of model [10] over 10,000 generations with $\Delta t$ = 1/12 years. Red lines denote +/- one standard deviation away from the (deterministic) equilibrium $\hat{p}$ as estimated from [A.13]. The blue line denotes $\hat{p}$ = 0.59.

The typical fluctuation size near $\hat{p}$ then has the simple form

$$\mathrm{Var}(P_t|P_0=\hat{p}) \approx \left(1-e^{-2R\prime(\hat{p})t}\right)\frac{F^2\mu(1-\mu)^2\hat{p}}{2N(\hat{p}(F-1)+1)^2(F-1-F\mu)} \quad \text{[A.12]}$$

$$= \left(1-e^{2R\prime(\hat{p})t}\right)\frac{\mu}{2N(F-1)}. \quad \text{[A.13]}$$

Figure A.1 gives an example with $F = 1.025$, $\mu = .01$, and $N = 10^4$ over 10000 iterations ($\approx$ 830 years) with initial conditions of $p_0 = \hat{p} = 0.59$. We observe that the time dependent factor in [A.13] converges to 0 very quickly, so that the variance converges to a constant. Instead of a long-time variance, we may rather be interested in a typical difference in $p_t$ over a single generation near $\hat{p}$. This quantity is simply estimated by

$$\sqrt{\mathrm{Var}(p_{t+1}|p_t=\hat{p})} = \sqrt{\frac{\mu\hat{p}}{FN(1-\mu)}} = 7.6\times 10^{-4}. \quad \text{[A.14]}$$

The distribution of $p_t$ in the discrete model [6] appears approximately normal, which can be explained by OU processes as well, as they have normal stationary distributions. Specifically, the stationary density for $P_t$ is distributed as

$$f_s \sim N\left(\hat{p}, \frac{\mu}{2N(F-1)}\right), \quad \text{[A.14]}$$

where $f \sim N(a, b^2)$ means that $f$ has a normal density with mean $a$ and variance $b^2$. From the model parameters above, we have an estimated mean $\hat{p} = 0.59$ and estimated standard deviation of $\sqrt{\frac{\mu}{2N(F-1)}} = 0.0045$.

**Table S1.** Populations at each stage of host reproduction and development. Here, $N$ is the initial female population, $I$ is the initial number of *Wolbachia*-positive females, $X_1 \sim \mathrm{Bin}(FpI, 1-\mu)$, and $X_2 \sim \mathrm{Bin}(F(1-p)I, 1-\mu)$.

| | $Wb^+$ Female $Wb^+$ Male | $Wb^+$ Female $Wb^-$ Male | $Wb^-$ Female $Wb^+$ Male | $Wb^-$ Female $Wb^-$ Male |
|---|---|---|---|---|
| **Matings** | $pI$ | $(1-p)I$ | $p(N-I)$ | $(1-p)(N-I)$ |
| **Ova** | $FpI$ | $F(1-p)I$ | $p(N-I)$ | $(1-p)(N-I)$ |
| **Ova after imperfect transmission** | $X_1$ | $X_2$ | $p(N-I)+FpI-X_1$ | $(1-p)(N-I)+F(1-p)I-X_2$ |
| **Adult offspring after CI** | $X_1$ | $X_2$ | $(1-s_h)(p(N-1)+FpI-X_1)$ | $(1-p)(N-I)+F(1-p)I-X_2$ |

**Table S2.** Regression analysis of the effects of model parameters on mean *Wolbachia* frequencies ($\bar{p}$).

| | | **Coefficient** | **t value** | ***P* value** |
|---|---|---|---|---|
| | **Host population size** | $-1.58 \times 10^{-8}$ | -6.09 | < 0.001 |
| **Imperfect maternal transmission ($\mu$)** | **$\mu$ value ($0.001 \leq \mu \leq 0.3$)** | $-2.24 \times 10^{1}$ | $-1.02 \times 10^{3}$ | < 0.001 |
| | **Low transmitters** | -2.43 | $-7.33 \times 10^{2}$ | < 0.001 |
| | **$\mu$ value * low transmitters** | 8.26 | $2.76 \times 10^{2}$ | < 0.001 |
| **Host fitness effects ($F$)** | **$F$ value ($1 \leq F \leq 1.5$)** | 3.77 | $4.01 \times 10^{2}$ | < 0.001 |
| | **Fluctuating $F$ ($CV = 0.01$)** | $1.30 \times 10^{-4}$ | $4.93 \times 10^{-2}$ | $9.61 \times 10^{-1}$ |
| | **Fluctuating $F$ ($CV = 0.1$)** | $9.68 \times 10^{-3}$ | 3.62 | < 0.001 |
| **Cytoplasmic incompatibility ($s_h$)** | **Weak CI ($s_h = 0.1$)** | $4.27 \times 10^{-1}$ | $1.63 \times 10^{2}$ | < 0.001 |
| | **Strong CI ($s_h = 0.45$)** | 1.92 | $6.82 \times 10^{2}$ | < 0.001 |

**Table S3.** Regression analysis of the effects of model parameters on the standard deviation of *Wolbachia* frequencies ($p_{SD}$).

| | | Coefficient | t value | *P* value |
|---|---|---|---|---|
| | **Host population size** | $-4.14 \times 10^{-7}$ | $-1.13 \times 10^{2}$ | < 0.001 |
| **Imperfect maternal transmission ($\mu$)** | **$\mu$ value ($0.001 \leq \mu \leq 0.3$)** | 8.51 | $4.03 \times 10^{2}$ | < 0.001 |
| | **Low transmitters** | 1.12 | $2.88 \times 10^{2}$ | < 0.001 |
| | **$\mu$ value * low transmitters** | -4.42 | $-1.39 \times 10^{2}$ | < 0.001 |
| **Host fitness effects (*F*)** | **$F$ value ($1 \leq F \leq 1.5$)** | -2.04 | $-1.68 \times 10^{2}$ | < 0.001 |
| | **Fluctuating *F* (*CV* = 0.01)** | $4.23 \times 10^{-1}$ | $1.01 \times 10^{2}$ | < 0.001 |
| | **Fluctuating *F* (*CV* = 0.1)** | 1.78 | $4.49 \times 10^{2}$ | < 0.001 |
| **Cytoplasmic incompatibility ($s_h$)** | **Weak CI ($s_h$ = 0.1)** | $-9.85 \times 10^{-2}$ | $-3.20 \times 10^{1}$ | < 0.001 |
| | **Strong CI ($s_h$ = 0.45)** | $-7.83 \times 10^{-1}$ | $-2.17 \times 10^{2}$ | < 0.001 |

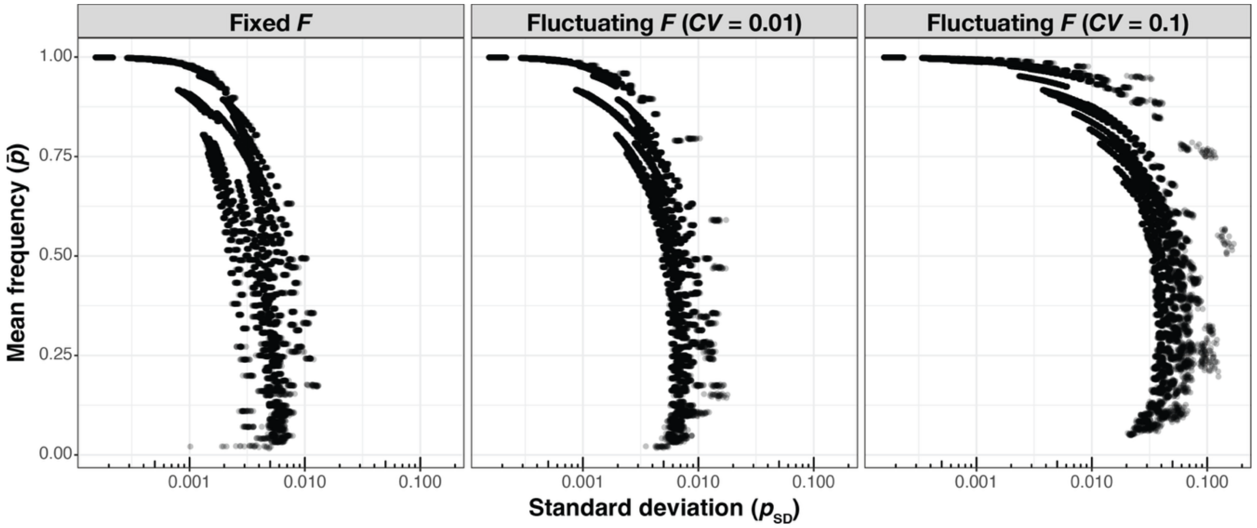


**Figure S1.** The relationship between mean infection frequency ($\bar{p}$) and the standard deviation ($p_{\text{SD}}$) in simulations with a host population of $N = 10^4$. Each individual point represents the $\bar{p}$ and $p_{\text{SD}}$ values from a single simulation. Datasets are separated based on whether fluctuating host effects ($CV = 0.01$ or $0.1$) are present.

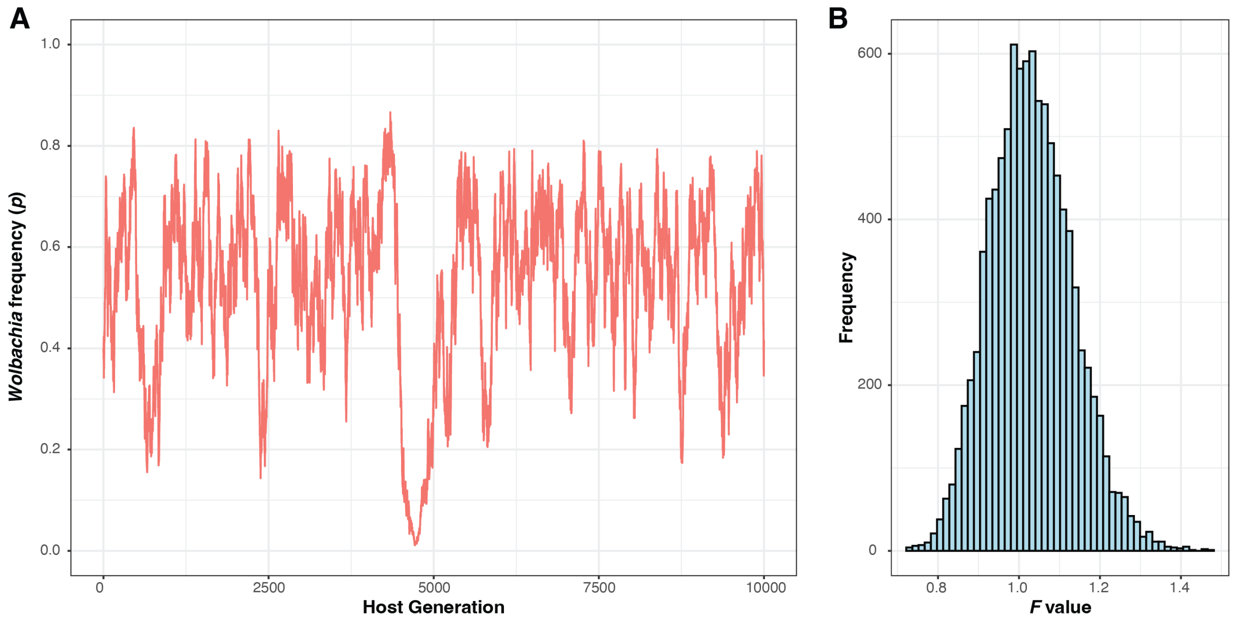


**Figure S2. (A)** For a host population size of $N = 10^4$ and fluctuating host fitness effects, the parameter values of $s_h = 0$, $F = 1.025$ ($CV = 0.1$), and $\mu = 0.01$ (without low transmitters) produced the largest average $p_{\mathrm{SD}}$ value of $\overline{p_{\mathrm{SD}}} = 0.144$ across 25 replicate simulations ($\bar{\bar{p}} = 0.540$). One of the 25 simulations is shown here as an example. **(B)** Histogram of $F$ values from 10,000 host generations for $F = 1.025$ ($CV = 0.1$). Strongly fluctuating $F$ values with a median slightly greater than one cause *Wolbachia* to alternate between favored ($F[1 - \mu] > 1$) and disfavored ($F[1 - \mu] < 1$) in the host population due to the large number of host generations with $F < 1$.

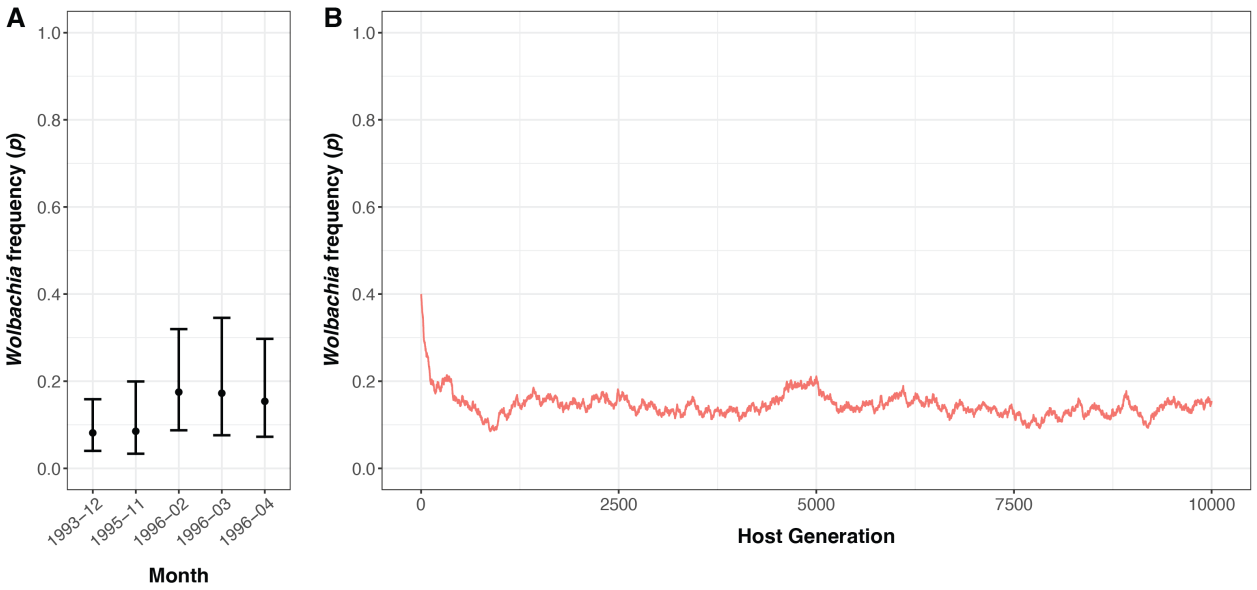


**Figure S3. (A)** Temporal fluctuations of *w*Mel frequencies ($p$) in *D. melanogaster* at Hastings in temperate eastern Australia. From the winter of 1993 to the spring of 1996, $p$ values exhibited minor fluctuations between a minimum of $p = 0.081$ (0.040, 0.159) and a maximum of $p = 0.175$ (0.087, 0.319). Data recreated from Hoffmann et al. (1998). **(B)** For a host population size of $N = 10^3$, the parameter values of $s_h = 0.1$, $F = 1.1$, and $\mu = 0.01$ (with low transmitters) produced an average of $\bar{\bar{p}} = 0.140$ and $\overline{p_{\mathrm{SD}}} = 0.020$ across 25 replicate simulations. One of the replicates is shown here as an example, where $p$ values fluctuated between a minimum of 0.085 and maximum of 0.211, which fall within the 95% binomial confidence intervals of the minimum and maximum *w*Mel frequencies observed at Hastings.

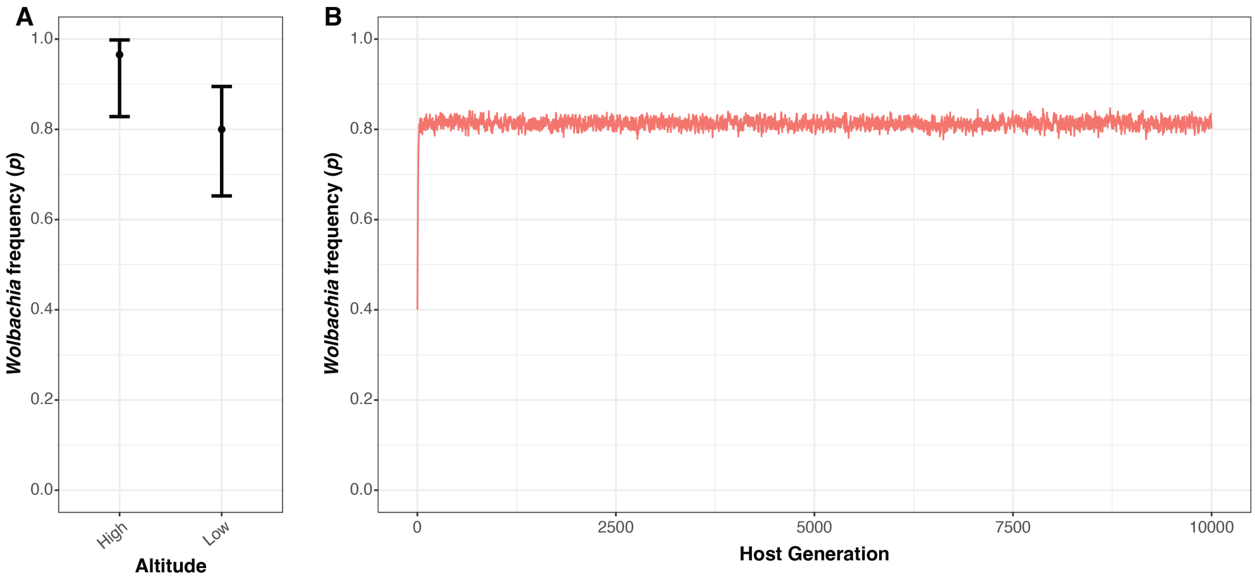


**Figure S4. (A)** Altitudinal variation of *w*Yak frequencies ($p$) in *D. yakuba* on the volcanic island of São Tomé off the coast of western Africa. In 2018, $p$ values differed between high ($p = 0.966$ [0.828, 0.998]) and low ($p = 0.800$ [0.652, 0.895]) altitude sites on the island. The high and low altitude sites are separated by 310 m of elevation and 2.4 km of distance. Data recreated from Hague et al. (2020). **(B)** For a host population size of $N = 10^3$, the parameter values of $s_h = 0.1$, $F = 1.225$, and $\mu = 0.05$ (no low transmitters) produced an average of $\bar{\bar{p}} = 0.813$ and $\overline{p_{\mathrm{SD}}} = 0.010$ across 25 replicate simulations. One of the replicates is shown here as an example, where $p$ values fluctuated between a minimum of 0.777 and maximum of 0.847, which fall within the 95% binomial confidence intervals of the high and low altitude *w*Yak frequencies observed at São Tomé in 2018. See Hague et al. (2020) for further details about the contribution of maternal transmission rates to altitudinal *w*Yak frequencies.

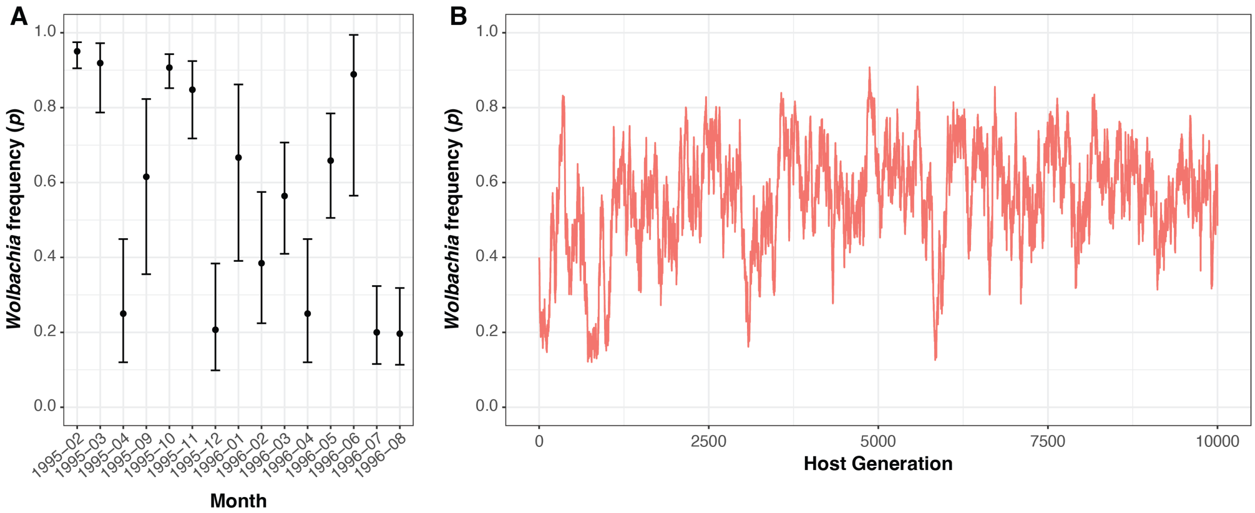


**Figure S5. (A)** Temporal fluctuations of *w*Mel frequencies ($p$) in *D. melanogaster* at Gold Coast in subtropical eastern Australia. From the winter of 1995 to the summer of 1996, $p$ values fluctuated between a minimum of $p = 0.196$ (0.113, 0.318) and a maximum of $p = 0.950$ (0.905, 0.975). Data recreated from Hoffmann et al. (1998). The authors reported no obvious seasonal effect on *w*Mel frequencies. **(B)** For a host population size of $N = 10^4$ and fluctuating host fitness effects, the parameter values of $s_h = 0$, $F = 1.025$ ($CV = 0.1$), and $\mu = 0.01$ (no low transmitters) produced an average of $\bar{\bar{p}} = 0.540$ and $\overline{p_{\mathrm{SD}}} = 0.144$ across 25 replicate simulations. One of the replicates is shown here as an example, where $p$ values fluctuated between a minimum of 0.121 and maximum of 0.908, which fall within the 95% binomial confidence intervals of the minimum and maximum *w*Mel frequencies observed at Gold Coast.

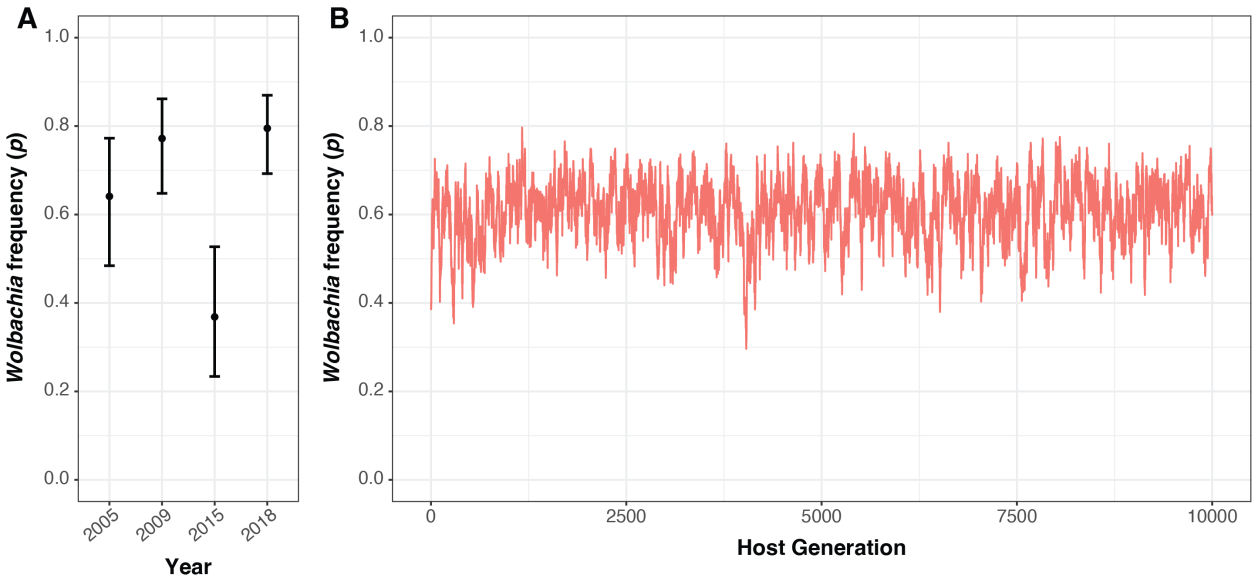


**Figure S6.** Temporal fluctuations of *w*San frequencies ($p$) in *D. santomea* on the volcanic island of São Tomé off the coast of western Africa. Over the course of 13 years, $p$ values fluctuated between a minimum of $p = 0.368$ (0.234, 0.527) and a maximum of $p = 0.795$ (0.692, 0.870). Data recreated from Hague et al. (2020). **(B)** For a host population size of $N = 10^4$ and fluctuating host fitness effects, the parameter values of $s_h = 0.1$, $F = 1.075$ ($CV = 0.1$), and $\mu = 0.05$ (no low transmitters) produced an average of $\bar{\bar{p}} = 0.606$ and $\overline{p_{\mathrm{SD}}} = 0.070$ across 25 replicate simulations. One of the replicates is shown here as an example, where $p$ values fluctuated between a minimum of 0.296 and maximum of 0.797, which fall within the 95% binomial confidence intervals of the minimum and maximum *w*San frequencies observed on São Tomé.
